# A small molecule inhibits *Neisseria gonorrhoeae* pilus formation

**DOI:** 10.64898/2026.11.19.767820

**Authors:** Kathleen R. Nicholson, Shaohui Yin, Kathryn A. Hayes, Saadman S. Ahmad, Brandon L. Jutras, Jennifer L. Edwards, Chi-Hao Luan, H Steven Seifert

## Abstract

*Neisseria gonorrhoeae* is becoming increasingly antimicrobial-resistant. *N. gonorrhoeae* requires the Type IV pilus (T4p) for pathogenesis, wherein the T4p promotes colonization through adherence to host cells and resistance to neutrophil killing. The pilus also is required for natural transformation and twitching motility. Collectively, these critical functions make the T4p an attractive drug target. Previously, we developed a high-throughput screen to identify compounds with activity against the zinc metalloprotease Mpg, which is involved in T4p biogenesis. Here, we identified a lead compound, SC144, that inhibits Mpg peptidoglycan-hydrolyzing activity in both *N. gonorrhoeae* and other bacteria encoding Mpg-like M23B metallopeptidase orthologs. Interestingly, SC144 targets Mpg’s carboxypeptidase and endopeptidase activities. Importantly, SC144 inhibited *N. gonorrhoeae* colonization and prevented its survival in two primary human cell culture models of infection without toxic effects on these host cells. This work provides strong evidence that SC144 is a promising lead compound with anti-virulence activity against *N. gonorrhoeae*.

**Author Summary:** Antimicrobial-resistant *Neisseria gonorrhoeae* infections are becoming prevalent. We targeted Mpg, a peptidoglycan-modifying enzyme important for pilus biogenesis, in an anti-virulence approach to identify novel antimicrobials. We show that the small molecule SC144 targets Mpg activity *in vitro* and alters peptidoglycan crosslinking similar to an Mpg mutant. SC144 also inhibits the enzymatic activity of several other M23 zinc metalloproteases. Mpg treatment reduces *N. gonorrhoeae* pilus formation, pilus-mediated adherence, and bacterial survival in cell culture models of infection. SC144 is a new compound that may be effective against gonorrhea and other bacterial infections.

## Introduction

*Neisseria gonorrhoeae* (the gonococcus) causes gonorrhea; however, a small, yet increasing subset of urogenital infections are caused by a specific clade of *Neisseria meningitidis* and are indistinguishable from gonorrhea [1,2]. *N. gonorrhoeae* infections are frequently asymptomatic, which increases transmission rates [1] and can lead to disseminated gonococcal infection with subsequent infectious arthritis and endocarditis [2,3]. Women with long-term gonococcal infections are at risk for severe reproductive health complications, such as infertility, ectopic pregnancies, pelvic inflammatory disease, and first-trimester abortion [4]. *N. gonorrhoeae* infection does not confer immunity to subsequent infections, and antibiotic resistance is rising [5–7]. Given its high transmission rate, lack of immediate symptoms, lack of a vaccine, and high rates of drug resistance, there is a critical need to identify novel therapeutics to target *N. gonorrhoeae*.

The Type IV pilus (T4p) is a major *N. gonorrhoeae* virulence factor. Gonococci isolated from infected human patients are usually piliated, whereas gonococci grown in the laboratory lose piliation during *in vitro* cultivation, suggesting piliation is selectively maintained during infection [8–12]. Pili mediate a variety of biological and pathogenic processes, including DNA uptake for transformation, twitching motility, adherence to host cells, biofilm formation, and resistance to killing by polymorphonuclear cells (PMNs) [13–16]. Thus, the T4p acts at multiple stages of infection, making it an attractive therapeutic target. However, pili, along with opacity and lipooligosaccharide genes, undergo antigenic and phase variation, both of which help evade host immunity and complicate the development of new treatments [5,17].

Previous studies have targeted the gonococcal pilus directly by screening for inhibitors of pilus function [18,19]. However, given the variable nature of pilin expression in *N. gonorrhoeae,* we reasoned that by indirectly inhibiting T4p functions, via non-phase variable enzymes, we could identify alternative therapeutic targets for future drug development that might offer a long-term solution to the growing problem of gonococcal antimicrobial resistance. Thus, we targeted a *non- phase variable* enzymatic component, Mpg (NGO_1686), which belongs to the M23B metallopeptidase family, as an alternative approach to inhibiting T4p function. M23B metallopeptidases participate in peptidoglycan remodeling in many bacterial species, including *Escherichia coli* and *Bacillus subtilis* [20–24]. Most of the bacterial M23B proteases target peptidoglycan peptide crosslinks [20–24]. Mpg has both carboxypeptidase and endopeptidase activities that are required for maximal T4p expression [16]. Loss of Mpg increases gonococcal sensitivity to PMN killing by decreasing pilus biogenesis [15], making it a viable drug that targets two essential colonization and pathogenesis activities.

We previously identified two natural compounds with activity against Mpg [25]. These compounds, punicalagin and chebulinic acid, decrease Mpg activity and T4p expression at 1 μM concentrations. Both compounds also reduced *N. gonorrhoeae* adherence to host cells and their survival in infection models with relevant host cell types [25]. However, these compounds have short half-lives in animals [26–28], making them unsuitable as therapeutic candidates. Here, we identified SC144, a quinoxalinhydrazide compound developed as an anticancer agent [29,30], as an inhibitor of Mpg’s protease activity and other M23B proteases. SC144 reduces T4p expression, and this anti-pilus activity, coupled with its previously established pharmacokinetics, supports its potential as a promising lead compound for gonorrhea treatment.

## Results

### SC144 binds to and inhibits Mpg protease activity

Using a fluorescence thermal shift (FTS) assay [25], we tested small-molecule binding to Mpg by thermal denaturation assays wherein fluorescence indicates Sypro Orange dye binding to hydrophobic regions of newly unfolded proteins. We identified SC144 (Fig 1A) as an Mpg-binding molecule in a screen of 37,760 compounds from five of the Northwestern High Throughput Analysis Laboratory’s compound libraries, including: 1) 1,280 molecules from the protein-protein interaction inhibitors collection, 2) 11,520 compounds from the kinase inhibitors collection, 3) 20,480 compounds from the diverse chemical structures collection, 4) 2,880 bioactive compounds, and 5) 1,600 compounds from the cell cycle/DNA damage collection [25]. We measured the FTS of Mpg upon treatment with 2.5 μM or 25 μM SC144 or a DMSO control (Fig 1B). Treatment with 2.5 μM SC144 did not alter the Mpg melting temperature (Tm) when compared to the DMSO control. However, treatment with 25 μM SC144 led to an ∼5 °C shift in the Tm.

**Fig 1.**
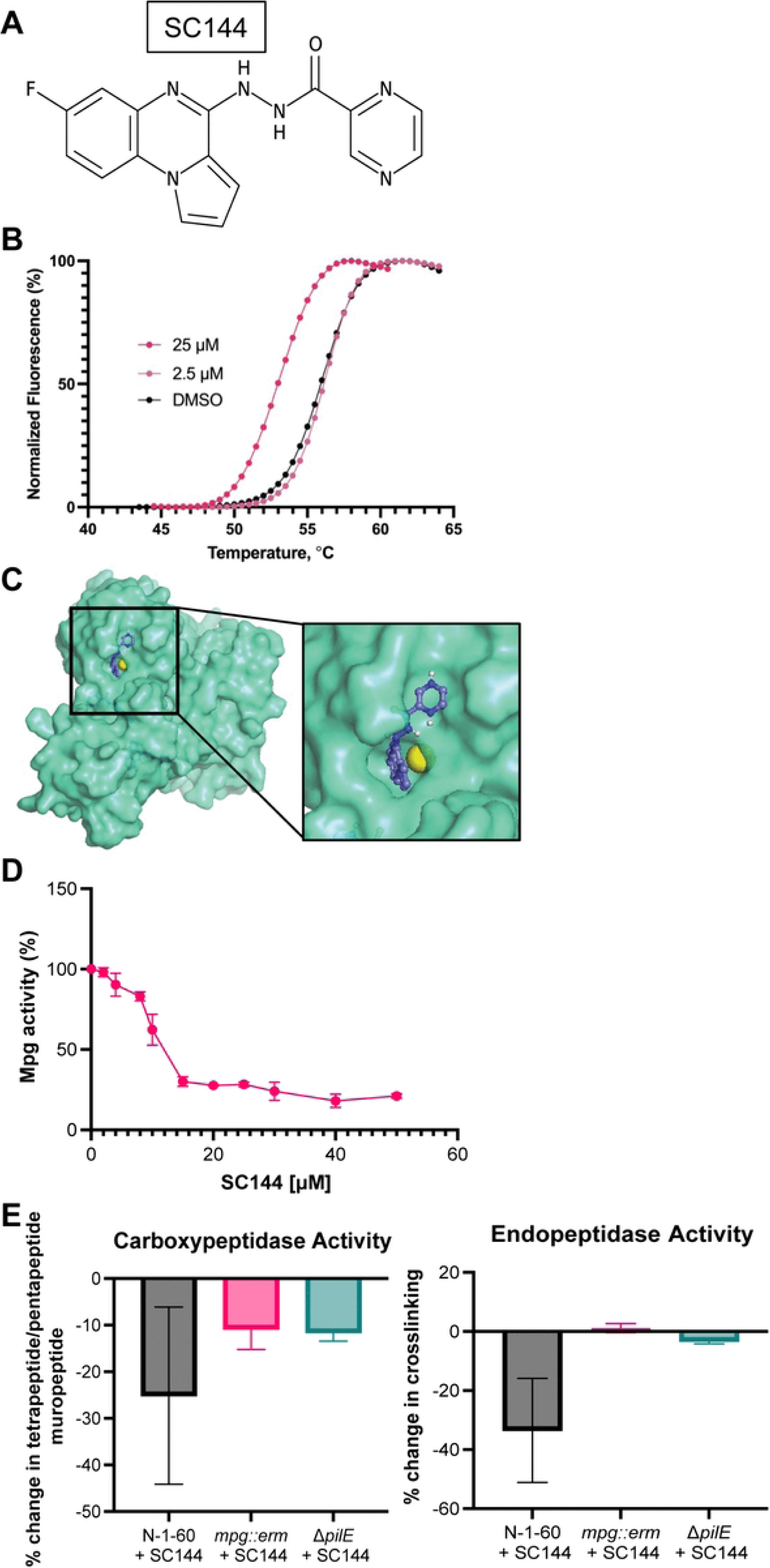
SC144 is a small molecule that binds to and has activity against Mpg. (A) Chemical structure of SC144. (B) Normalized compound melting curves with DMSO control. (C) Space filling model of molecular docking analysis of Mpg and SC144 using Autodock and Alphafold predicted SC144 binding in the Mpg active site. Molecular graphs were generated in Pymol with color scheme: Mpg in green/cyan and zinc in yellow, SC144: carbon in blue, and hydrogen in white. (D) Peptidoglycan (PG) dye release assay with an Mpg dose response. 0.5 – 50 μM of SC144 reacted with 1 μM Mpg. (E) LCMS analysis of 0.75 μM SC144-treated gonococci. **Left**, relative loss of endopeptidase activity in SC144 treated samples. Endopeptidase activity was calculated based on the total peak area of crosslinked muropeptides divided by the total peak area of the sample. Percent change was calculated between treated and untreated samples for each genetic background. The data represented are three biological replicates. **Right**, relative loss in carboxypeptidase activity in SC144 treated samples. Carboxypeptidase activity was calculated by dividing the total peak area of tetrapeptide PG monomers by the total peak area of pentapeptide PG monomers. Percent change was calculated between treated and untreated samples for each genetic background. The data represented are three biological replicates.

Molecular docking with AutoDock Vina software predicted that SC144 binds to the Mpg active site (Fig 1C), which is evolutionarily conserved (S1 Fig). Since Mpg is an M23B metallopeptidase with peptidoglycan-hydrolyzing activity, we used an *in vitro* peptidoglycan dye release assay [31] to quantify the ability of SC144 to inhibit Mpg protein activity (Fig 1D). With 12 μM of SC144, Mpg (1 µM) activity decreased by 50%. Interestingly, 50 μM SC144 reduced Mpg activity to 20% of its baseline, with no further reduction at higher amounts. These data show that SC144 targets and inhibits Mpg activity *in vitro*.

Mpg exhibits both carboxypeptidase and endopeptidase activity [16]. To determine if SC144 targets both activities *in vivo*, we measured peptidoglycan fragments from treated and untreated gonococci using LCMS analysis (S2 Fig) [32,33]. We calculated carboxypeptidase activity by quantifying the ratio of tetrapeptide to pentapeptide muropeptide monomers, where a higher ratio indicates more carboxypeptidase activity. We then compared this ratio in cells treated with 0.75 μM SC144 compared to untreated cells. We used a *N. gonorrhoeae* FA1090 derivative, N-1-60, with mutations in the *pilE* guanine quadruplex sequence and the *pilC2* sequence to prevent antigenic and phase variation, ensuring consistent pilus expression as our parental strain [34]. SC144 treatment of the N-1-60 strain showed a decrease in carboxypeptidase activity, consistent with reduced Mpg enzymatic activity (Fig 1E, left). Despite a downward trend, this difference was not statistically significant compared to the *mpg::erm* and Δ*pilE* strains, due to a large spread in replicate data. The *mpg::erm* and Δ*pilE* strains showed slight decreases in carboxypeptidase activity similar to each other, consistent with a loss of Mpg activity through genetic deletion or loss of the T4p. We then determined endopeptidase activity by calculating the percentage of crosslinked muropeptides in each sample. Like carboxypeptidase activity, endopeptidase activity decreased, albeit not statistically significantly, in the N-1-60 parental strain under SC144 treatment, consistent with a loss of Mpg activity (Fig 1E, right). The *mpg::erm* and Δ*pilE* strains showed no appreciable change in endopeptidase activity. Collectively, these data suggest SC144 inhibits Mpg enzymatic activity *in vivo*.

### SC144 reduces gonococcal piliation

Mutation of Mpg, or treatment with natural compounds that inhibit Mpg, reduces *N. gonorrhoeae* T4p expression [16,25]. Because SC144 targets and inhibits Mpg activity *in vitro*, similar to other natural compounds that target Mpg [25], we determined whether SC144 treatments alters *N. gonorrhoeae* piliation. We assayed SC144’s effect on pilus-dependent colony morphology in the N-1-60 strain and the *mpg::erm* mutant. Piliated *N. gonorrhoeae* have small, domed, and dark colony morphologies, whereas nonpiliated *N. gonorrhoeae* colony morphologies are larger, flat, and without dark shading when observed with a stereo microscope (Fig 2A) [8,35]. Growth of piliated *N. gonorrhoeae* in medium with 0.5 μM SC144 produced colonies that were larger, flatter, and with lighter coloration, similar to an *mpg* mutant, while with 1 μM SC144, we noted slight decreases in colony size (Fig 2A). We used immunogold transmission electron microscopy (immunogold-TEM) to visualize pili on the surface of individual N-1-60 gonococci with a carboxy-terminus Myc-tagged *pilE* allele [34] in the presence or absence of SC144. Without SC144, gonococci produce normal T4p fibers as evidenced by gold nanoparticle labeling (Fig 2B). In the presence of 1 μM SC144, fewer and shorter T4p were present on the cell surface similar to an *mpg* mutant [15]. These data show that 1 μM SC144 treatment reduced T4p elaboration.

**Fig 2.**
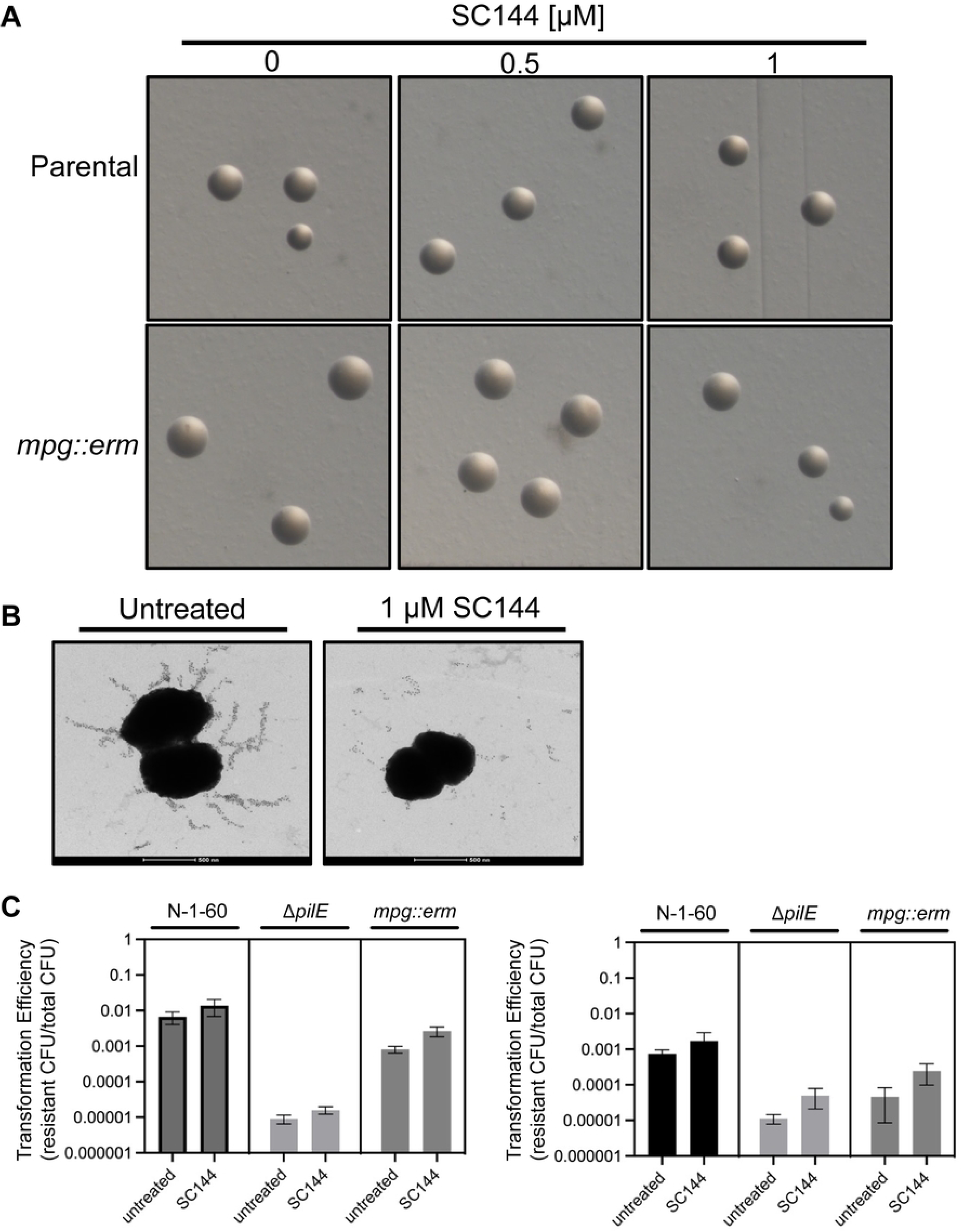
Treatment with SC144 reduces *N. gonorrhoeae* piliation. (A) Colony morphology of *N. gonorrhoeae* grown with 0.5 or 1 μM of SC144. Colonies were imaged after 22 hours of growth. Piliated colonies appear smaller and darker than nonpiliated colonies. (B) Immunogold-TEM of pili on gonococci treated with 1μM SC144. Assays were performed in a strain containing a *pilE-myc* allele, and pilus expression was detected using an α-Myc primary antibody and a 5 nm gold-conjugated secondary antibody. 18500X.

### Treatment with SC144 reduces adherence and survival in infection cell culture models

Gonococcal T4p mediate adherence to host cells and tissues during colonization, the first step of infection [36]. Because SC144 treatment decreases piliation, we wanted to determine whether SC144 treatment influenced the adherence of two *N. gonorrhoeae* strains, MS11 and FA1090, to primary male urethral cells (UECs) and cervical (Pex) epithelial cells [37,38]. We first confirmed that SC144 was not toxic to UECs, Pex or HeLa cells (S3 Fig). There was a statistically significant decrease in adherence of both MS11 (to approximately 20%) and FA1090 (to approximately 17%) to Pex cells with 0.5 μM SC144 compared to the untreated control (Fig 3A). With 1 μM and 2 μM SC144, both MS11 and FA1090 adherence decreased to approximately 4% and 0.7%, respectively. Interestingly, at 4 μM SC144, no adherent bacteria were detected (Fig 3A). Adherence to UEC cells showed similar trends with both MS11 and FA1090 showing significantly decreased adherence (to approximately 18% and 10%, respectively) with 0.5 μM SC144, approximately 4% adherence at 1 μM, 1% adherence at 2 μM, and no detectable adherent bacteria at 4 μM (Fig 3B). Collectively, these data show that micromolar concentrations of SC144 prevent *N. gonorrhoeae* adherence to host cells.

**Fig 3.**
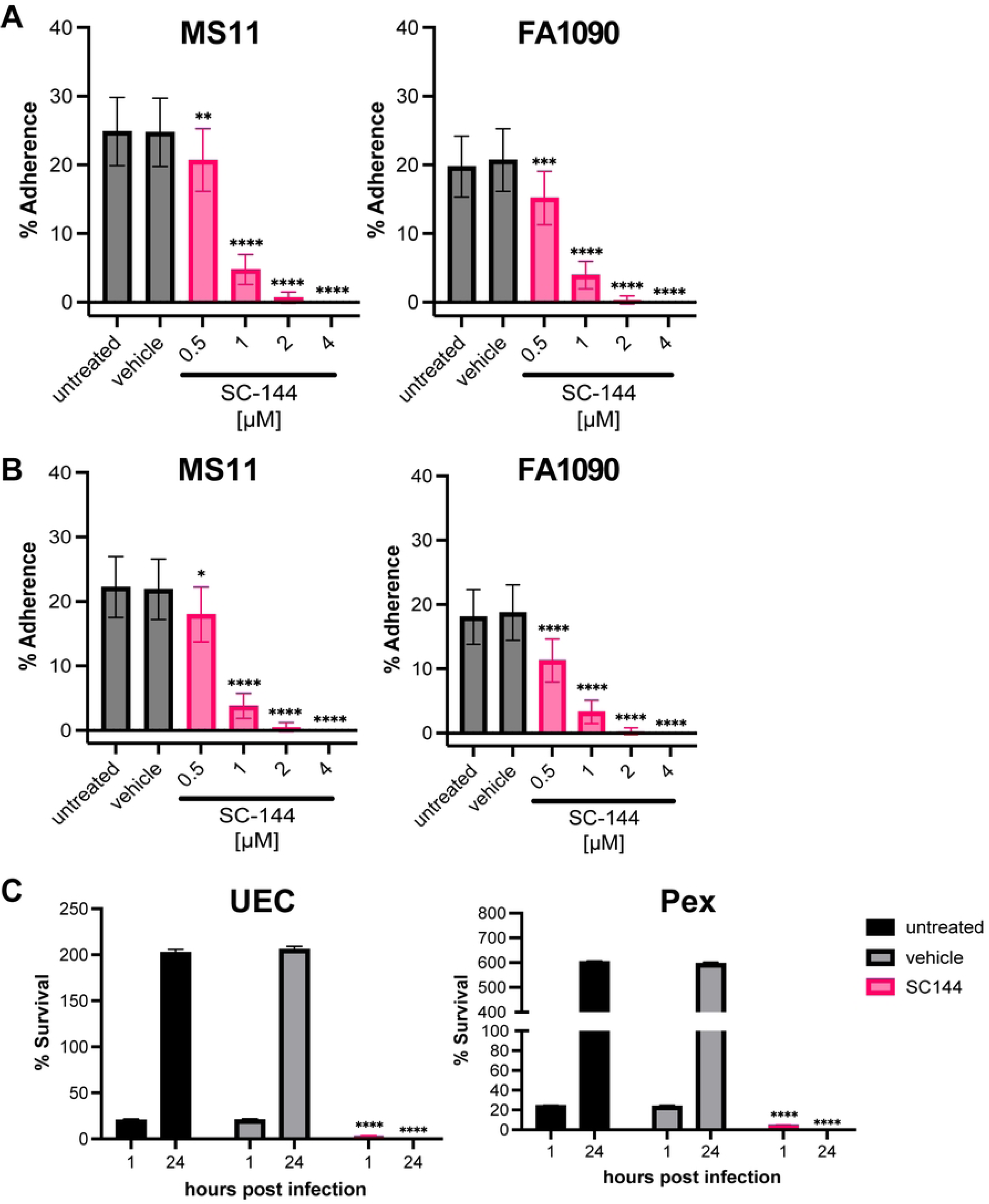
SC144 treatment decreases adherence of *N. gonorrhoeae* to UEC and Pex cells. *N. gonorrhoeae* MS11 or FA1090 that were pre-treated (1h) without or with 0.1% DMSO (vehicle), or (0.5, 1, 2 or 4 µM) SC-144 was then used to infect (A) Pex cells or (B) UECs (MOI=100). Following a 1h infection, the medium was removed, cells were rinsed, lysed, and serial dilutions were plated. Compound was maintained in the medium throughout the infection. The percentage of *N. gonorrhoeae* associated with host cells was determined as a function of each respective inoculum. A nonparametric ANOVA was used to determine statistical significance. ****, p<0.0001; ***, p<0.001; **, p<0.01; *, p<0.05. (C) N. gonorrhoeae strain MS11 pre-treated (1h) without or with 0.1% DMSO (vehicle) or 1 µM SC-144 was used to infect **Left.** UECs or **Right.** Pex cells (MOI=100) for 1h or 24h. At each time point, the infection medium was removed, cells were rinsed, lysed, and serial dilutions were plated. Compound was maintained in the medium throughout the infection. The percentage of *N. gonorrhoeae* associated with host cells or that survived treatment was determined as a function of the inoculum. A nonparametric ANOVA was used to determine statistical significance. ****, p<0.0001.

Pili play a critical role in adherence to mucosal epithelial cells, which also initiates gonococcal invasion and intracellular survival within host cells during mucosal infection. In addition to measuring initial adherence, we measured the effect of SC144 treatment on gonococcal survival after 24 hours of infection of UECs and Pex cells [37,38]. We used the MS11 strain to quantify both adherence and survival in the presence of 1 μM SC144. Untreated bacteria served as a positive control. In the absence of SC144, MS11 adhered to and survived within UECs, resulting in bacterial replication by 24h post-infection with an approximate 200% increase in survival compared to the initial inoculum. The 0.1% DMSO vehicle control behaved similarly to untreated bacteria. Upon the addition of 1 μM SC144, adherence was significantly reduced compared to the untreated control, and no bacteria were detected after 24 hours (Fig 3C). In Pex cells, untreated bacteria and vehicle-treated bacteria adhered and showed approximately 600% survival at the 24-hour time point (Fig 3C). Treatment with SC144 significantly reduced initial adherence, and no bacteria were recovered after 24 hours of infection. These data indicate that treatment with 1 μM SC144 is sufficient to reduce the initial adherence of gonococci to relevant host cells and to prevent their survival and/or replication by 24 hours of infection.

### SC144 shows inhibitory activity against Mpg orthologs

M23B zinc metallopeptidases have many established roles in both Gram-positive and Gram-negative bacterial species, and many M23 proteases have roles in modifying peptidoglycan [20–24]. To determine whether SC144 has activity against other M23B enzymes, we purified M23 metallopeptidases from a set of Gram-negative and Gram-positive bacteria. We measured the peptidoglycan-hydrolyzing activity of purified Gram-negative M23 metallopeptidases with or without 50 μM SC144. The purified *N. meningitidis* Mpg ortholog showed statistically significantly reduced enzymatic activity in the presence of SC144, while treatment with SC144 nearly ablated the activity of the purified *Pseudomonas aeruginosa* and *Acinetobacter baumanii* M23 metallopeptidases (Fig 4A). The activity of the *Vibrio cholerae* and *Escherichia coli* M23 metallopeptidases was also significantly reduced under SC144 treatment (Fig 4A). Similarly, we found SC144 inhibited the peptidoglycan-hydrolyzing activity of *S. aureus* LytM by approximately 50% and also the peptidoglycan-hydrolyzing activity of a *S. aureus* glycine-glycine endopeptidase (Fig 4B). These results strongly suggest that SC144 could have biological activity against other Gram negative and Gram positive organisms,

**Fig 4.**
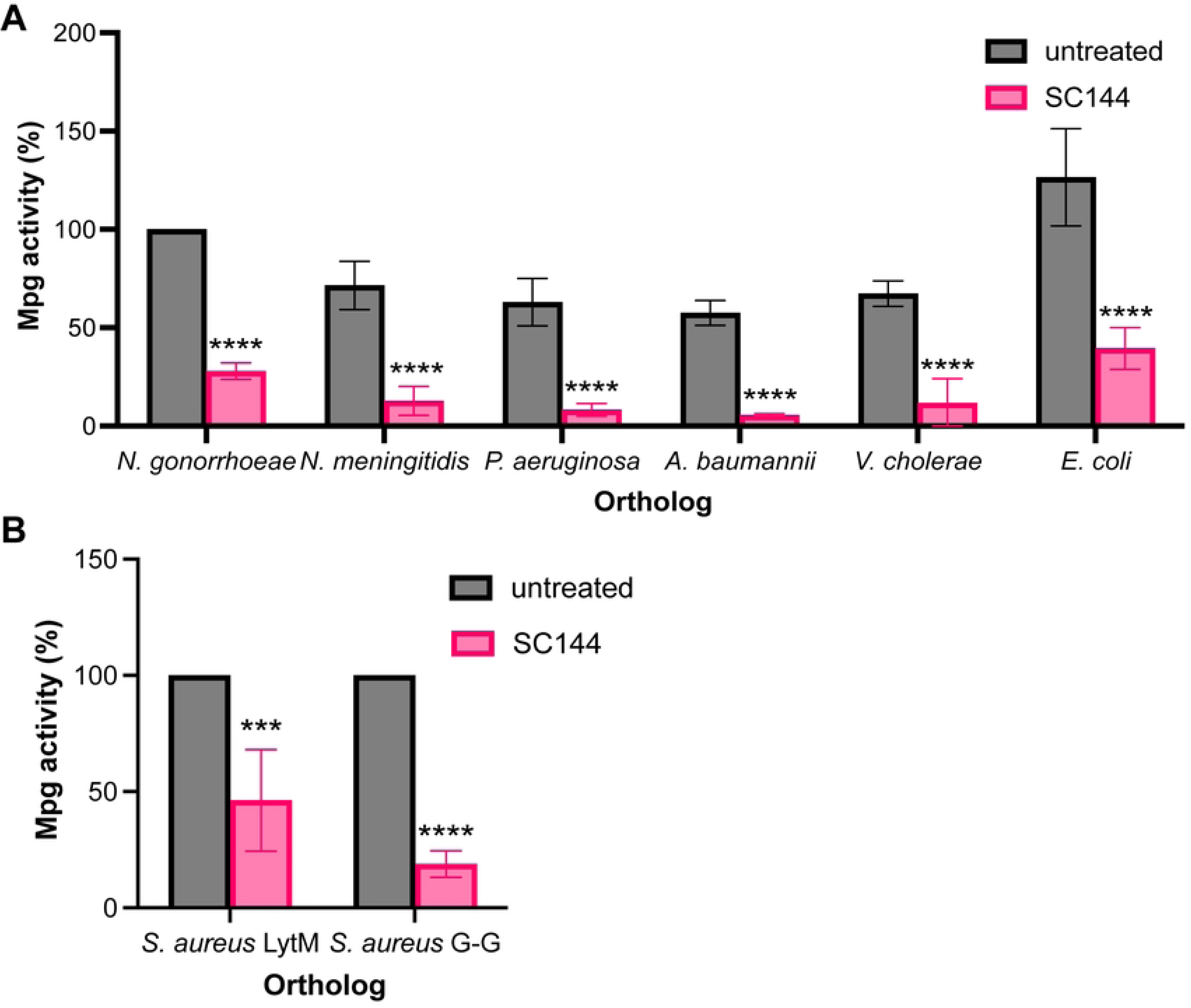
SC144 inhibits Mpg orthologs. (A) Mpg orthologs from Gram-negative species*: N. gonorrhoeae* FA1090; *N. meningitidis* MC58; *P. aeruginosa* PAO1; *A. baumannii* Ab2*; V. cholerae* O395 VC1; *E. c*oli K12 MG1655. Each reaction contained 0.5 μM Mpg, 50 μM SC144, and 10 mg/ml RBB-labeled gonococcal PG. Percent activity is represented relative to untreated Mpg activity. Statistical analysis was performed using a two-way ANOVA with Šidák’s multiple comparisons. Significance is shown relative to untreated Mpg from each bacterial ortholog (****, p<0.0001). (B) Mpg orthologs from Gram-positive *S. aureus*. Each reaction contained 0.5 μM Mpg, 50 μM SC144, and 10 mg/ml RBB-labeled gonococcal PG. Percent activity is represented relative to untreated Mpg activity. Statistical analysis was performed using a two-way ANOVA with Šidák’s multiple comparisons. Significance is shown relative to untreated (****, p<0.0001; ***, p = .0.0001).

## Discussion

We targeted the enzymatic activity of the Mpg protease as part of an anti-virulence strategy. Using this approach, we identified SC144 as a novel inhibitor of Mpg activity in the bacterial cell and *N. gonorrhoeae* piliation. SC144 was previously identified as a novel quinoxalinhydrazide compound with potent anticancer activity [29,30]. It acts as a glycoprotein-130 (GP130) inhibitor that targets the interleukin-6 and Stat3 signaling axis in mammalian cells [30]. Multiple studies have examined the behavior of SC144 in animal models, including mice and pigs [39]. SC144 shows oral activity in mouse models of ovarian cancer and specificity for targeting cancer cells, with limited toxicity against nonmalignant cells [29,30]. In translational large-animal porcine models, SC144 showed activity as an injectable for treating right ventricular dysfunction (RVD), which was alleviated by inhibiting GP130 signaling [39]. These previous studies suggest that SC144, or analogous compounds, could be used in humans although there are not human studies published to date.

We chose to target the Mpg M23B zinc metallopeptidase, as it is required for full pilus elaboration and is not targeted by short-term diversity generating mechanisms like phase or antigenic variation. Our evolutionary analyses indicate Mpg active site residues (H295, D299, H373, H375) [16], at the predicted SC144 binding site, are >90% conserved [40] amongst orthologs. Outside the conserved active site histidine residues, the larger region encompassing these residues is also highly conserved. Thus, while evolutionary pressure for drug resistance may increase, these residues are less prone to mutation on shorter evolutionary timescales. Using peptidoglycan hydrolyzing assays, we showed SC144 broadly inhibits Mpg enzymatic activity *in vitro*. Additionally, we indirectly measured carboxypeptidase and endopeptidase activity in gonococci treated with SC144. We found that in the parental strain expressing Mpg, carboxypeptidase and endopeptidase activity trended downward in the presence of SC144. We conclude that SC144 is active on Mpg *in vitro* and *in vivo* and likely targets both enzymatic activities.

Further, we showed treating gonococci with SC144 at concentrations below 1 μM targeted Mpg enzymatic activity, resulting in a concomitant reduction in T4p biogenesis. Most gonorrhea infections occur at the genital mucosa; therefore, we used primary, human male urethral and female cervical epithelial cells to assess gonococcal T4p-mediated adherence. We showed SC144 was not toxic to either host cell type at concentrations as high as 8 μM, consistent with previously published studies [29,30]. During infection, treatment with 1 μM SC144 significantly reduced pilus-mediated adherence and prevented gonococcal survival after 24 hours of infection. Notably, SC144 cured (100% clearance, no CFUs detected) *N. gonorrhoeae* MS11 urethral and cervical cell infections. Ultimately, SC144’s pilus-targeting activity prevents the initial step of host colonization in two *N. gonorrhoeae* strains. Moreover, we also showed that SC144 retains *in vitro* activity against M23B metallopeptidases from *P. aeruginosa*, *E. coli*, *V. cholerae,* and *S. aureus*, indicating it may have broader activity in other bacteria.

This study identified a novel small molecule with therapeutic potential in gonorrhea treatment. SC144 exhibited anti-virulence activity in gonococci, targeting a major virulence factor (i.e., pilus) through a non-phase variable, enzymatic component of the T4p apparatus. SC144’s potent activity in cell culture models of gonorrhea infection, and limited toxicity in small- and large-animal models, make it a promising candidate for future infection and pharmacokinetic studies involving *N. gonorrhoeae*.

## Materials and Methods

### Bacterial Strains and Media

Gonococcal strain MS11 and derivatives of the FA1090 strain, N-1-60 [34], were used herein. In the N-1-60 strain, the guanine quadruplex site upstream of the *pilE* gene is mutated to prevent antigenic variation, and the *pilC2* allele is locked in an “on” conformation to ensure expression of pili [34]. Gonococcal strains were grown on Gc medium base (GCB; Difco) (36.25g/L), agar (1.25g/L), Kellogg supplement I (22.2mM glucose, 0.68mM glutamine, 0.45 mM cocarboxylase) and II [1.23 mM Fe(NO_3_)_3_] [9,41]. We added compounds to the medium before solidification at the concentrations indicated in the text. *N. gonorrhoeae* was grown on GCB agar with or without compounds at 37 °C and 5% CO_2_ for 22 hours. S1 Table lists detailed strain information.

### Generation of DNA Constructs

We generated all DNA constructs using oligonucleotide primers purchased from Integrated DNA Technologies (IDT, Coralville, IA). We verified all resulting constructs by targeted Sanger sequencing (GeneWiz). All constructs and oligonucleotides are listed in S2 Table and S3 Table, respectively..

### Protein Expression Constructs

We PCR-amplified the *mpg* gene and its orthologs (without the predicted signal sequence) from chromosomal DNA of the noted bacterial species using a forward primer with an NheI site and a reverse primer with a HindIII site, respectively. The PCR fragment was digested by NheI/HindIII restriction enzymes and cloned into an NheI/HindIII digested pET28a overexpression vector to yield a 6X His-tagged Mpg.

### Mpg and M23 Metallopeptidase Expression and Purification

To purify the 6X His-Mpg or M23 metallopeptidase orthologs, *E. coli* BL21(DE3) cells (Novagen) harboring the pET28a construct containing the *mpg* or M23 metallopeptidase ortholog gene were inoculated into 4 ml LB medium containing kanamycin (50 μg/ml), grown overnight at 37 °C. These cultures were then 50X diluted into fresh medium containing kanamycin (35 μg/ml), grown for another 2-3 hours (OD_600_ 0.4-0.5), and induced by addition of 1mM IPTG for 4 hours at 37 °C. The cells were pelleted by centrifugation, resuspended in cold 1X His-Bind Buffer (Kit; Novagen) at 20 ml per 100 ml volume of cell culture medium (Novagen), and sonicated on ice using a Vibra-Cell VC250 with a microtip (Sonics & Materials) at 35% duty cycle for a total of 3-4min in pulsed mode, followed by centrifugation (15,000 x g) for 20 min to clarify the protein extract. The extract was purified on Novagen Ni-NTA His-Bind resin by column chromatography according to the manufacturer’s protocol (TB054 Rev. F0106). The eluate, containing purified His-Mpg orthologs, was dialyzed against sodium acetate buffer (20 mM, pH 4.5, 10% glycerol) and concentrated using a 10 kDa MWCO Amicon® Ultra Centrifugal Filter at 4 °C.

### Library and Screening Using a Fluorescence Thermal Shift Assay

37,760 molecules were screened using fluorescence thermal shift (FTS) assays from several Northwestern High Throughput Analysis Laboratory’s compound library: 1,280 molecules from the protein-protein interaction inhibitors collection, 11,520 from the kinase inhibitors collection (ChemDiv Inc., CA), 20,480 diverse chemical structures collection (ChemBridge Corp., CA), 2,880 bioactive compounds (TargetMol Chemicals Inc., MA), and 1600 compounds from the cell cycle/DNA damage collection (MedChemExpress USA, NJ).

Recombinant Mpg showed consistent performance in FTS assays, with a standard deviation of 0.12 °C across 32 replicates and a T_m_ of 57 °C. The assay was conducted in 384-well microplates, with Mpg (1 μM or 0.04 μg/μL) premixed with a 5X concentration of Sypro-Orange in HEPES buffer (20 mM HEPES, 150 mM NaCl, pH 7.5). 10 or 5 μL of the protein-dye mixture was dispensed into each well, and 10 to 50 nanoliters of compounds (10-50 μM, prepared from 10 mM stock solutions in DMSO) were added using an Echo550 acoustic transfer robot (Labcyte, CA). After sealing with optical tape, centrifuging, and mixing, we performed a thermal scan from 20 °C to 90 °C at 0.5 °C/min, collecting fluorescence data on a CFX384 real-time PCR machine (Bio-Rad Laboratories).

### Peptidoglycan Isolation and Dye Labeling

Peptidoglycan (PG) was purified from *N. gonorrhoeae* and labeled with RBB (Remazol brilliant blue, Sigma), as described previously [15]. Briefly, overnight FA1090 cells from 40 plates were swabbed in 20 ml cold PBS buffer and centrifuged at 4500 rpm for 10 min at 4 °C. The pellet was resuspended in 10 ml PBS buffer, boiled on a hot plate and adjusted to 4% SDS by adding 10 ml 8% SDS in PBS, and kept boiling for 2 hours. After boiling twice, the pellet was collected by ultracentrifugation (28000 RPM, SI TW32, 25 °C, 30 min), and washed 3x with 1X PBS buffer. The sacculi pellet was resuspended in 10 ml PBS containing Pronase (1 mg/ml), DNase and RNase (50 mg/ml each), incubated for 2 hours to overnight at 37 C, and ultracentrifuged to remove the supernatant. The pellet was boiled again in 20 ml 4% SDS for 2-4 hours, washed 3x with PBS, resuspended in 10-20 ml 0.25 N NaOH containing 20 mM RBB, and incubated at 37°C overnight. The reaction was neutralized by adding an equal volume of 0.25 N HCl, and ultracentrifuged. The RBB-PG pellet was then thoroughly washed with 20 ml of sterilized water 7-8 times by ultracentrifugation.

### Dye Release Assay for PG Hydrolysis

10 mg/ml of RBB-PG, 1 μM purified Mpg, and compounds at indicated concentrations were incubated for four hours at 37 °C in a total of 15 ml of assay buffer (25 mM Tris-HCl (pH 7.4), 100 mM NaCl, 0.5% Triton X-100). We terminated the reaction by heating at 95 °C for 5 minutes, then spun at 14,000 rpm for 10 minutes in a microcentrifuge at room temperature. We collected 10 ml of supernatant and measured its absorbance at 595 nm with a NanoDrop spectrophotometer. We have performed this assay with varying concentrations of both Triton X-100 or Tween-20, and 0.5% Triton X-100 is optimal for this assay.

### SC144 treatment for PG purification and LCMS analysis

*N. gonorrhoeae* strains were struck from -80 °C stocks and grown on GCB agar overnight at 37 °C under 5% CO_2_. Bacteria were swabbed (polyester swab) into 1X PBS and adjusted to 0.5X McFarland standard (∼10^8^ CFU/ml). 100 μl of this culture was plated on a square petri dish containing GCB agar with or without 0.75 μM SC144 (MedChemExpress) compound. Plates were incubated overnight at 37 °C under 5% CO_2_. Bacteria from five plates (per sample) were harvested in 1X PBS using polyester swabs followed by a 1X PBS wash over the entire plate to gather remaining bacteria. Cells were pelleted by centrifugation (4000 rpm x 10 min).

### PG purification for LCMS analysis

Pelleted *N. gonorrhoeae* was resuspended in PBS and added dropwise to boiling 10% sodium dodecyl sulfate (SDS), for a final SDS concentration of 5%. Cell suspensions were boiled for 1 hour and insoluble sacculi were harvested via ultracentrifugation (50,000 rpm, 1 hour, 35 °C). Contaminating SDS was removed from the sacculi by four rounds of washes with warm ultrapure water before being treated with 0.5 mg/mL chymotrypsin in 1 mM CaCl_2_ and 2mM HCl overnight. Proteases were denatured with 0.5% SDS for 30 min and sacculi were harvested using methods described above. Purified sacculi were digested into muropeptide fragments by overnight incubation with 60 U/mL of mutanolysin. The next morning an additional 60 U/mL of mutanolysin was added and allowed to digest an additional 4-6 hours. The mutanolysin was heat inactivated for 10 min at 100 °C and the sample was centrifuged at 22,000 x *g* for 30 min to pellet any insoluble material. Soluble muropeptides in the supernatant were concentrated via lyophilization. Dried purified muropeptides were resuspended in 150 µL of saturated sodium borate buffer, pH 9.25. Next, a 50 µL aliquot of 100 mg/mL NaBH_4_ was added in a dropwise manner. The reaction was quenched after 1 h with approximately 10 µL of formic acid to a final pH of 3, determined with colorimetric pH paper. The samples were then snap frozen and lyophilized. The dried muropeptides were resuspended in H_2_O:MeCN (200 µL, 9:1, v:v) containing 0.1% formic acid and sonicated in a water bath for 10 min followed by centrifugation (13,000 g, 4°C, 10 min). 180 µL of the supernatant was transferred to an LCMS vial for analysis.

### LCMS analysis

Analyses were performed as previously published [32,33] on a Shimadzu LCMS9030 QToF system coupled to a LC-40D X3 UPLC, a SIL-40C X3 autosampler (10 °C), and a CTO-40C column oven (40 °C). Borohydride reduced mutanolysin-digested muropeptides were separated on a Waters BEH C18 column (2.1×100 mm, 130 Å pore size, 1.7 µm particle size), at an initial flow rate of 0.4 ml/min. Solvent A was water and Solvent B was MeOH, with both solvents containing 0.1% formic acid (v/v). The initial solvent condition was 99:1 (A:B) for 3 min. This was followed by stepwise linear gradients first to 8% Solvent B (12 min), then to 20% Solvent B (24 min) and an adjusted flow rate of 0.3 ml/min, and finally 5% Solvent B (25 min), was followed by a hold at 5% B until 30 min, at which the flow rate was returned to initial conditions. The column was then re-equilibrated to 1% B (1 min to 1% B followed by a 5 min re-equilibration time). The reduced muropeptides were subjected to electrospray ionization in positive ion mode using data dependent acquisition. The interface voltage was 4.0 kV at 300 °C with a desolvation temperature of 526 °C and a DL temperature of 250 °C. Nebulizing, heating, and drying gas flows were 2, 10, and 10 L/min, respectively. The m/z range for the MS scan was 300 to 2000.

### Imaging of Pilus-Dependent Colony Morphology and Pilus Detection

For colony morphology imaging, *N. gonorrhoeae* colonies grown for 22 hours on solid medium were observed and photos were taken using a Nikon SMZ-10A stereomicroscope and a Nikon digital-site camera.

To analyze the effect of the compound on the piliation of the gonococcal strain on solid medium, immunoelectron microscopy was performed as described previously [34]. Briefly, Formvar/carbon-coated copper grids were used to lift cells directly from 18 h old colonies and fixed for 15 min by adding a drop (17 μl) of 0.2% glutaraldehyde and 4% paraformaldehyde in Dulbecco’s PBS (DPBS; Fisher) to the grids. The grids were washed 3 times with 1% bovine serum albumin (BSA; Sigma) in DPBS and blocked in 0.1% gelatin (Aurion, Inc.) in DPBS for 30 minutes. The grids were washed once with 1% BSA in DPBS and incubated with a 1:10 dilution of rabbit anti-c-Myc antibody (Sigma) for 1 h. Grids were washed three times with 1% BSA in DPBS and incubated with 0.1% gelatin in PBS for 30 min. The grids were washed once with BSA in DPBS and incubated with goat anti-rabbit immunoglobulin G antibody conjugated to 12-nm gold particles (1:20 dilution; Jackson ImmunoResearch) for 1 h. Grids were washed five times in water for 3 minutes each. The grids were negatively stained with 1% uranyl acetate for 1 min. All washes and incubations were 17 μl and performed at room temperature. The liquid on the grids after each step was carefully wicked away using a Whatman paper. Grids were viewed using a FEI Tecnai Spirit G2 transmission electron microscope (TEM).

### UEC and Pex Cell cultures

Primary male urethral cells (UECs) and cervical (Pex) epithelial cells were procured from de-identified human tissues, as described [37,38]. In brief, this process involves culturing epithelial cells from dissected urethral or cervical tissue. Urethral tissue was obtained from the National Disease Research Interchange (Philadelphia, PA, USA), whereas cervical tissue was obtained from the Cooperative Human Tissue Network (Columbus, OH, USA). We maintained tissue explants and primary epithelial cells in human urethral epithelial cell medium (Cell Applications, San Diego, CA, USA) for UECs or defined keratinocyte serum-free medium (dk-SFM; Gibco, Grand Island, NY, USA) for Pex cells. The use of these tissues does not constitute human subjects research, as determined by the Institutional Review Board at the Abigail Wexner Research Institute at Nationwide Children’s Hospital.

### UEC and Pex Host Cell Viability Assays

To determine if SC-144 exerted a cytotoxic effect toward uninfected UECs or Pex cells, cells were incubated for 24h with 0.1% DMSO (vehicle); 0.5, 1, 2, 4, or 8 µM of SC144; or they were left untreated. We determined UEC and Pex cell viability fluorometrically using the Cell Viability Assay Kit (Fluorometric-Blue), according to the manufacturer’s instructions (Abcam, Waltham, MA USA). We performed assays in duplicate on three separate occasions. Data were adjusted for background after which fluorescence recorded for each condition was normalized to fluorescence recorded for untreated cells. A paired Student’s t-test (GraphPad, GraphPad Software, San Diego, CA, USA) was used to determine the statistical significance of data obtained.

### UEC and Pex Cell Infection Assays

Infection assays were performed by pre-incubating (1h, 37 °C) bacteria with medium alone or with medium supplemented with 0.1% DMSO (vehicle) or 0.5, 1, 2, or 4 µM SC-144, as noted, before infection of each host cell type. For association assays, bacteria were incubated (37 °C) with host cells for 1h following pre-treatment with each drug, whereas survival assays comprised of a 24h incubation using the MS11 strain. The FA1090 strain becomes cytotoxic in primary cells during 24h infections, making this strain unsuitable for these experiments. DMSO (vehicle control) or SC-144 was maintained in the medium throughout the course of infection. For both assays, host cell monolayers were rinsed following infection and lysed. Serial dilutions of the host cell lysates were plated, and viable gonococci were enumerated by counting CFUs after a 48h incubation (37 °C, 5% CO_2_). We determined the percentage of *N. gonorrhoeae* that associated with, or that survived within, host cells as a function of the inoculum used to initiate each infection. All assays were performed in triplicate on 3 separate occasions using a multiplicity of infection of 100. We used a non-parametric ANOVA to determine the statistical significance of bacterial association or survival (GraphPad Prism).

### HeLa Cell Culture

HeLa cells were cultured in Dulbecco’s Modified Eagle’s Medium (DMEM) (Gibco, Dublin, Ireland) with 10% heat-inactivated fetal bovine serum (FBS) (VWR, Radnor, PA). Cells were washed with 1X sterile phosphate-buffered saline (PBS) (Gibco) and harvested using 0.25% trypsin-EDTA digestion (Gibco). All experiments were performed at 37 °C with 5% CO_2_.

### HeLa MTS Assays

HeLa cells were seeded in 100 μl DMEM plus 10% FBS per well at 2×10^4^ cells/well in a 96 well plate (Grenier bio-one, Germany) and allowed to grow for 24h. Media was removed, cells were washed, and 80 μl of fresh media was added. Compounds were added in a 20 μl volume to final concentrations of 1, 2, 3, or 5 μΜ in technical triplicate. Treatment was allowed to proceed for 24h before media was removed and cells were washed three times with 1X PBS. 100 μl of 1X PBS was added to each well. 20 μl of MTS reagent (CellTiter 96 AQ_ueous_ One Solution Cell Proliferation Assay, Promega) was added to each well. Cells were incubated with MTS reagent for 1h at 37°C under 5% CO_2_ before absorbance (A_490_) was read on a Spectramax M5 spectrophotometer. Assays were performed in biological triplicate.

### HeLa Live/Dead Viability/Cytotoxicity Assays

HeLa cells were seeded to 96 well plates and cultured as described above. Compounds, in 20 μl volumes to final concentrations of 1, 2, 3, or 5 μM, were added to the HeLa cells (in 80 µl of culture medium/well). Treatments, in technical triplicate, were allowed to proceed for 24 h before the medium was removed, and the cells were washed three times with 1X PBS. To assay viability/cytotoxicity, PBS was aspirated, and 100 μl of an ethidium homodimer-1 (EthD-1; 4 μM) and Calcein AM (2 μM) solution (Live/Dead Viability/Cytotoxicity Kit; Life Technologies, Carlsbad, CA) in 1X PBS was added. Cells were incubated for 30 minutes before fluorescence was measured (Calcein AM: 494 nm excitation/517 nm emission; EthD-1: 528 nm excitation/617 nm emission) on a Spectramax M5 plate reader. Assays were performed in biological triplicate.

### Statistical Analysis

All statistical analyses were performed in GraphPad Prism 11 version 11.0.2. Statistical significance was determined by performing an ordinary one-way ANOVA followed by a Dunnett’s multiple comparisons test or two-way ANOVA followed by a Šidák’s multiple comparisons test. P-values for individual experiments are noted in figure legends and in the text. All experiments were conducted on at least two biological replicates, each with technical triplicates where available.

## Acknowledgments

This work was supported by the Northwestern University High Throughput Analysis Laboratory and the Northwestern University Center for Advanced Microscopy. Technical advice was provided by Gary Schlitz of the Northwestern University Chemistry Department. We thank the Cooperative Human Tissue Network/Human Tissue Resource Network, Columbus, OH, USA, which is funded by the National Cancer Institute, for cervical tissue specimens. We thank Joseph Dillard from the University of Wisconsin-Madison for the useful discussion.

## Data Availability

The authors confirm that all supporting data and protocols have been provided within the article and its supplemental material. The raw data generated and analyzed during this study are openly available in the PRISM repository at https://doi.org/10.18131/3da89-tb834.

## Supporting Information Captions

**S1 Fig. ConSurf-DB analysis of Mpg.** (A) ConSurf-DB [40] residue conservation mapping on crystal structure of Mpg (PDB: 6MUK) illustrating evolutionary conservation at each residue. Scale from Variable (1) to Conserved (9). Yellow indicates insufficient data. (B) Mpg amino acid conservation from ConSurf-DB. This sequence does not include the signal sequence.

**S2 Fig. Representative chromatograms of peptidoglycan isolated from parental and SC144 treated strains.** Highlighted peaks are monomeric peptidoglycan fragments which were used in the analysis. Each chromatogram is representative of multiple biological replicates (n=3 N-1-60, N-1-60+SC144, *mpg::erm*, and *mpg::erm*+SC144; n=2 Δ*pilE*, Δ*pilE*+SC144). Strains were treated with 0.75 μM SC144.

**S3 Fig. SC144 acid does not decrease HeLa cell metabolic activity or lead to cytotoxicity.** (A) MTS assay measuring metabolic activity of HeLa cells grown in the presence of increasing concentrations (1 μM-5 μM) SC144. Doxorubicin serves as a positive control for decreased cellular metabolism. (B) Live/Dead cytotoxicity assay. HeLa cells were grown with increasing concentrations of SC144. Live cells were quantified using calcein-AM (1 μM) staining. Dead cells were quantified using Ethidium homodimer (EthD-1; 2 μM). Calcein-AM was measured at an excitation of 494 nm and emission of 517 nm. EthD-1 was measured at an excitation of 528 nm and emission of 617 nm. Percent live cells was calculated relative to untreated cells stained with Calcein-AM. Percent dead cells was calculated relative to 0.01% saponin-treated cells stained with EthD-1 or EthD-1. Statistical analysis was performed using a two-way ANOVA with multiple comparisons to doxorubicin at indicated concentrations. (C) UEC or Pex cells incubated for 24h with noted agent before rinsing and addition of fluorescence assay reagent using the Abcam Viability Assay Kit. Cells were incubated for 1h at 37°C before reading 530/25 excitation; 645/40 emission. Increased fluorescence corresponds to increased cell viability. A paired student’s t-test was used to determine statistical significance. **, p<0.01; *, p<0.05.

**S1 Table. Strains used in this study.**

**S2 Table. Plasmids used in this study.**

**S3 Table. Oligonucleotides used in this study.**

## Literature Cited

1. 1. WHO guidelines for the treatment of Neisseria gonorrhoeae. World Health Organization; 2016. 55 p.

2. Agabawi S, Tran V, McEachern J, Walkty A. Gonococcal meningitis: An unusual presentation of disseminated gonococcal infection. J Assoc Med Microbiol Infect Dis Can. 2019 Jun;4(2):116–20. doi:10.3138/jammi.2018-0043 PubMed PMID: 36337739.

3. Masi AT, Eisenstein BI. Disseminated gonococcal infection (DGI) and gonococcal arthritis (GCA): II. Clinical manifestations, diagnosis, complications, treatment, and prevention. Semin Arthritis Rheum. 1981 Feb;10(3):173–97. doi:10.1016/s0049-0172(81)80002-9 PubMed PMID: 6785887.

4. Hook EW, Holmes KK. Gonococcal infections. Ann Intern Med. 1985 Feb;102(2):229–43. doi:10.7326/0003-4819-102-2-229 PubMed PMID: 3917638.

5. Schmidt KA, Schneider H, Lindstrom JA, Boslego JW, Warren RA, Van de Verg L, et al. Experimental gonococcal urethritis and reinfection with homologous gonococci in male volunteers. Sex Transm Dis. 2001 Oct;28(10):555–64. doi:10.1097/00007435-200110000-00001 PubMed PMID: 11689753.

6. Berçot B, Caméléna F, Mérimèche M, Jacobsson S, Sbaa G, Mainardis M, et al. Ceftriaxone-resistant, multidrug-resistant Neisseria gonorrhoeae with a novel mosaic penA-237.001 gene, France, June 2022. Euro Surveill. 2022 Dec;27(50). doi:10.2807/1560-7917.ES.2022.27.50.2200899 PubMed PMID: 36695466.

7. Ohnishi M, Golparian D, Shimuta K, Saika T, Hoshina S, Iwasaku K, et al. Is Neisseria gonorrhoeae initiating a future era of untreatable gonorrhea?: detailed characterization of the first strain with high-level resistance to ceftriaxone. Antimicrob Agents Chemother. 2011 Jul;55(7):3538–45. doi:10.1128/AAC.00325-11 PubMed PMID: 21576437.

8. Kellogg DS, Peacock WL, Deacon WE, Brown L, Pirkle DI. Neisseria Gonorrhoeae. I. VIRULENCE GENETICALLY LINKED TO CLONAL VARIATION. J Bacteriol. 1963 Jun;85(6):1274–9. doi:10.1128/jb.85.6.1274-1279.1963 PubMed PMID: 14047217.

9. Swanson J, Robbins K, Barrera O, Corwin D, Boslego J, Ciak J, et al. Gonococcal pilin variants in experimental gonorrhea. J Exp Med. 1987 May 1;165(5):1344–57. doi:10.1084/jem.165.5.1344 PubMed PMID: 3106555.

10. Hamrick TS, Dempsey JAF, Cohen MS, Cannon JG. Antigenic variation of gonococcal pilin expression in vivo: analysis of the strain FA1090 pilin repertoire and identification of the pilS gene copies recombining with pilE during experimental human infection. Microbiology (Reading). 2001 Apr;147(Pt 4):839–49. doi:10.1099/00221287-147-4-839 PubMed PMID: 11283280.

11. Jerse AE, Cohen MS, Drown PM, Whicker LG, Isbey SF, Seifert HS, et al. Multiple gonococcal opacity proteins are expressed during experimental urethral infection in the male. J Exp Med. 1994 Mar 1;179(3):911–20. doi:10.1084/jem.179.3.911 PubMed PMID: 8113683.

12. Boiko I, Metaane S, Seifert HS. Publicly Available Neisseria Gonorrhoeae Genomes Predominantly Represent In Vitro-Derived Nonpiliated Variants. J Infect Dis. 2026 Feb 18;233(2):e311–21. doi:10.1093/infdis/jiaf557 PubMed PMID: 41334759.

13. Sparling PF. Genetic transformation of Neisseria gonorrhoeae to streptomycin resistance. J Bacteriol. 1966 Nov;92(5):1364–71. doi:10.1128/jb.92.5.1364-1371.1966 PubMed PMID: 4958881.

14. Merz AJ, So M. Interactions of pathogenic neisseriae with epithelial cell membranes. Annu Rev Cell Dev Biol. 2000;16:423–57. doi:10.1146/annurev.cellbio.16.1.423 PubMed PMID: 11031243.

15. Stohl EA, Dale EM, Criss AK, Seifert HS. Neisseria gonorrhoeae metalloprotease NGO1686 is required for full piliation, and piliation is required for resistance to H2O2- and neutrophil-mediated killing. mBio. 2013 Jul 9;4(4). doi:10.1128/mBio.00399-13 PubMed PMID: 23839218.

16. Stohl EA, Chan YA, Hackett KT, Kohler PL, Dillard JP, Seifert HS. Neisseria gonorrhoeae virulence factor NG1686 is a bifunctional M23B family metallopeptidase that influences resistance to hydrogen peroxide and colony morphology. J Biol Chem. 2012 Mar 30;287(14):11222–33. doi:10.1074/jbc.M111.338830 PubMed PMID: 22334697.

17. Wielert I, Kraus-Römer S, Volkmann TE, Craig L, Higgins PG, Maier B. Pilin antigenic variants impact gonococcal lifestyle and antibiotic tolerance by modulating interbacterial forces. PLoS Biol. 2025 Jan 30;23(1):e3003022. doi:10.1371/journal.pbio.3003022

18. Aubey F, Corre JP, Kong Y, Xu X, Obino D, Goussard S, et al. Inhibitors of the *Neisseria meningitidis* PilF ATPase provoke type IV pilus disassembly. Proceedings of the National Academy of Sciences. 2019 Apr 23;116(17):8481–6. doi:10.1073/pnas.1817757116

19. Denis K, Le Bris M, Le Guennec L, Barnier JP, Faure C, Gouge A, et al. Targeting Type IV pili as an antivirulence strategy against invasive meningococcal disease. Nat Microbiol. 2019 Mar 25;4(6):972–84. doi:10.1038/s41564-019-0395-8

20. Boneca IG. The role of peptidoglycan in pathogenesis. Curr Opin Microbiol. 2005 Feb;8(1):46–53. doi:10.1016/j.mib.2004.12.008 PubMed PMID: 15694856.

21. Goley ED, Comolli LR, Fero KE, Downing KH, Shapiro L. DipM links peptidoglycan remodelling to outer membrane organization in Caulobacter. Mol Microbiol. 2010 Jul 1;77(1):56–73. doi:10.1111/j.1365-2958.2010.07222.x PubMed PMID: 20497504.

22. Möll A, Schlimpert S, Briegel A, Jensen GJ, Thanbichler M. DipM, a new factor required for peptidoglycan remodelling during cell division in Caulobacter crescentus. Mol Microbiol. 2010 Jul 1;77(1):90–107. doi:10.1111/j.1365-2958.2010.07224.x PubMed PMID: 20497502.

23. Poggio S, Takacs CN, Vollmer W, Jacobs-Wagner C. A protein critical for cell constriction in the Gram-negative bacterium Caulobacter crescentus localizes at the division site through its peptidoglycan-binding LysM domains. Mol Microbiol. 2010 Jul 1;77(1):74–89. doi:10.1111/j.1365-2958.2010.07223.x PubMed PMID: 20497503.

24. Uehara T, Dinh T, Bernhardt TG. LytM-domain factors are required for daughter cell separation and rapid ampicillin-induced lysis in Escherichia coli. J Bacteriol. 2009 Aug;191(16):5094–107. doi:10.1128/JB.00505-09 PubMed PMID: 19525345.

25. Nicholson KR, Yin S, Edwards JL, Luan CH, Seifert HS. Natural compounds target the M23B zinc metallopeptidase Mpg to modulate Neisseria gonorrhoeae Type IV pilus expression. mBio. 2025 Apr 9;16(4):e0402724. doi:10.1128/mbio.04027-24 PubMed PMID: 39998224.

26. Dhingra AK, Chopra B, Grewal AS, Guarve K. Pharmacological properties of Chebulinic acid and related ellagitannins from nature: An emerging contemporary bioactive entity. Pharmacological Research - Modern Chinese Medicine. 2022 Dec;5:100163. doi:10.1016/j.prmcm.2022.100163

27. Lu Y, Yan H, Teng S, Yang X. A liquid chromatography-tandem mass spectrometry method for preclinical pharmacokinetics and tissue distribution of hydrolyzable tannins chebulinic acid and chebulagic acid in rats. Biomed Chromatogr. 2019 Mar;33(3):e4425. doi:10.1002/bmc.4425 PubMed PMID: 30397912.

28. Xu J, Cao K, Liu X, Zhao L, Feng Z, Liu J. Punicalagin Regulates Signaling Pathways in Inflammation-Associated Chronic Diseases. Antioxidants (Basel). 2021 Dec 24;11(1). doi:10.3390/antiox11010029 PubMed PMID: 35052533.

29. Grande F, Aiello F, Garofalo A, Neamati N. Identification and Preclinical Evaluation of SC144, a Novel Pyrroloquinoxaline Derivative with Broad-Spectrum Anticancer Activity. Mini Rev Med Chem. 2016;16(8):644–50. doi:10.2174/138955751608160307175649 PubMed PMID: 26955877.

30. Xu S, Grande F, Garofalo A, Neamati N. Discovery of a novel orally active small-molecule gp130 inhibitor for the treatment of ovarian cancer. Mol Cancer Ther. 2013 Jun;12(6):937–49. doi:10.1158/1535-7163.MCT-12-1082 PubMed PMID: 23536726.

31. Stohl EA, Lenz JD, Dillard JP, Seifert HS. The Gonococcal NlpD Protein Facilitates Cell Separation by Activating Peptidoglycan Cleavage by AmiC. J Bacteriol. 2016 Feb 15;198(4):615–22. doi:10.1128/JB.00540-15

32. Ahmad SS, Ebohon O, McClune ME, Trimble RN, Kellogg CN, Booth CJ, et al. Peptidoglycan architecture dictates protein interactions, tissue tropism, and arthritis in the Lyme disease spirochete Borrelia burgdorferi. PLoS Pathog. 2026 Jan;22(1):e1013849. doi:10.1371/journal.ppat.1013849 PubMed PMID: 41557633.

33. DeHart TG, Kushelman MR, Hildreth SB, Helm RF, Jutras BL. The unusual cell wall of the Lyme disease spirochaete Borrelia burgdorferi is shaped by a tick sugar. Nat Microbiol. 2021 Nov 24;6(12):1583–92. doi:10.1038/s41564-021-01003-w

34. Hu LI, Yin S, Ozer EA, Sewell L, Rehman S, Garnett JA, et al. Discovery of a New Neisseria gonorrhoeae Type IV Pilus Assembly Factor, TfpC. mBio. 2020 Oct 27;11(5). doi:10.1128/mBio.02528-20 PubMed PMID: 33109763.

35. Swanson J, Kraus SJ, Gotschlich EC. Studies on gonococcus infection. I. Pili and zones of adhesion: their relation to gonococcal growth patterns. J Exp Med. 1971 Oct 1;134(4):886–906. doi:10.1084/jem.134.4.886 PubMed PMID: 4106489.

36. Higashi DL, Lee SW, Snyder A, Weyand NJ, Bakke A, So M. Dynamics of Neisseria gonorrhoeae attachment: microcolony development, cortical plaque formation, and cytoprotection. Infect Immun. 2007 Oct;75(10):4743–53. doi:10.1128/IAI.00687-07 PubMed PMID: 17682045.

37. Harvey HA, Ketterer MR, Preston A, Lubaroff D, Williams R, Apicella MA. Ultrastructural analysis of primary human urethral epithelial cell cultures infected with Neisseria gonorrhoeae. Infect Immun. 1997 Jun;65(6):2420–7. doi:10.1128/iai.65.6.2420-2427.1997 PubMed PMID: 9169783.

38. Edwards JL, Shao JQ, Ault KA, Apicella MA. Neisseria gonorrhoeae elicits membrane ruffling and cytoskeletal rearrangements upon infection of primary human endocervical and ectocervical cells. Infect Immun. 2000 Sep;68(9):5354–63. doi:10.1128/IAI.68.9.5354-5363.2000 PubMed PMID: 10948165.

39. Mendelson JB, Sternbach JD, Kim M, Moon RA, Hartweck LM, Clark SR, et al. GP130 Antagonism Enhances Porcine RV Function. Circ Res. 2025 Oct 24;137(10):1231–51. doi:10.1161/CIRCRESAHA.125.326336 PubMed PMID: 41031411.

40. Ben Chorin A, Masrati G, Kessel A, Narunsky A, Sprinzak J, Lahav S, et al. ConSurf-DB: An accessible repository for the evolutionary conservation patterns of the majority of PDB proteins. Protein Sci. 2020 Jan;29(1):258–67. doi:10.1002/pro.3779 PubMed PMID: 31702846.

41. Dillard JP. Genetic Manipulation of Neisseria gonorrhoeae. Curr Protoc Microbiol. 2011 Nov;Chapter 4:Unit4A.2. doi:10.1002/9780471729259.mc04a02s23 PubMed PMID: 22045584.

